# Cooperation of dominant oncogenes with regulatory germline variants shapes clinical outcomes in childhood cancer

**DOI:** 10.1101/506659

**Authors:** Julian Musa, Florencia Cidre-Aranaz, Marie-Ming Aynaud, Martin F. Orth, Olivier Mirabeau, Mor Varon, Sandrine Grossetête, Didier Surdez, Shunya Ohmura, Julia S. Gerke, Aruna Marchetto, Marlene Dallmayer, Michaela C. Baldauf, Moritz Gartlgruber, Frank Westermann, Stefanie Stein, Tilman L. B. Hölting, Maximilian M. L. Knott, Giuseppina Sannino, Jing Li, Laura Romero-Pérez, Wolfgang Hartmann, Uta Dirksen, Melissa Gymrek, Nathaniel D. Anderson, Adam Shlien, Barak Rotblat, Thomas Kirchner, Olivier Delattre, Thomas G. P. Grünewald

## Abstract

Deciphering principles of inter-individual tumor heterogeneity is essential for refinement of personalized anti-cancer therapy. Unlike cancers of adulthood, pediatric malignancies including Ewing sarcoma (EwS) feature a striking paucity of somatic alterations except for pathognomonic driver-mutations that cannot explain overt variations in clinical outcome.

Here we demonstrate in the EwS model how cooperation of a dominant oncogene and regulatory variants determine tumor growth, patient survival and drug response.

We show that binding of the oncogenic EWSR1-FLI1 fusion transcription factor to a polymorphic enhancer-like DNA element controls expression of the transcription factor MYBL2, whose high expression promotes poor patient outcome via activation of pro-proliferative signatures. Analysis of paired germline and tumor whole-genome sequencing data revealed that regulatory variability at this locus is inherited via the germline. CRISPR-mediated interference with this regulatory element almost abolished *MYBL2* transcription, and MYBL2 knockdown decreased cell proliferation, cell survival and tumorigenicity of EwS cells. Combined RNA- and ChIP-seq analyses as well as functional experiments and clinical data identified *CCNF, BIRC5* and *AURKB* as direct MYBL2 targets and critical mediators of its phenotype. In drug-response experiments, high MYBL2 levels sensitized EwS cells for inhibition of its activating cyclin dependent kinase CDK2 *in vitro* and *in vivo,* suggesting MYBL2 as a predictive biomarker for targeted anti-CDK2-therapy.

Collectively, our findings establish cooperation of somatic mutations and regulatory germline variants as a major determinant of tumor progression and indicate the importance of integrating the regulatory genome in the process of developing new diagnostic and/or therapeutic strategies to fully harness the potential of precision medicine.

## MAIN TEXT

The advent of high-throughput ‘omics’-technologies in clinical oncology enabled assignment of patients to targeted therapies based on somatic mutations in the protein coding genome^1^. However, many childhood cancers including Ewing sarcoma (EwS) – a highly aggressive bone-associated cancer – hardly exhibit any recurrent genetic alteration other than pathognomonic and uniformly expressed driver-mutations^2,3^. Yet, these tumors show substantial inter-individual heterogeneity concerning clinical behavior and treatment response, which cannot be solely explained by their few (epi-)genetic alterations^3–5^.

However, recent studies in model organisms suggested that the effects of dominant oncogenes may depend on inter-individual variations in the regulatory genome^6^. Thus, we hypothesized that oncogenic cooperation of driver-mutations with specific regulatory germline variants may explain inter-individual diversity in clinical outcomes of oligo-mutated cancers.

We explored this possibility in EwS, which constitutes a genuine model to study such cooperation for several reasons: First, it is characterized by a relatively simple, nearly diploid genome with a single driver-mutation resulting from chromosomal rearrangements fusing the *EWSR1* gene to various members of the ETS family of transcription factors (in 85% FLI1)^2,3,7,8^. Second, EWSR1-FLI1 steers ~40% of its target genes by binding DNA at GGAA-microsatellites, which are thereby converted into potent enhancers^9–11^. Third, the enhancer activity of EWSR1-FLI1-bound GGAA-microsatellites strongly depends on the inter-individually variable number of consecutive GGAA-repeats^9,10,12^. Together, these characteristics provide an ideal framework to analyze how cooperation of a dominant oncogene (here EWSR1-FLI1) with polymorphic regulatory elements (here GGAA-microsatellites) influences the expression of disease-promoting genes that could explain clinical diversity in oligo-mutated cancers.

To identify such candidate genes with high clinical relevance, we crossed two datasets. The first comprised gene expression microarrays of A673 EwS cells harboring a doxycycline (DOX)-inducible shRNA against *EWSR1-FLI1* (A673/TR/shEF1) profiled with/without addition of DOX (**Supplementary Table 1**). The second comprised 166 transcriptomes of primary EwS with matched clinical annotation (**Supplementary Table 2**). We calculated for each gene represented in both datasets the fold change (FC) of its expression after DOX-induced EWSR1-FLI1 knockdown in A673/TR/shEF1 cells and the significance levels for association with overall survival (OS) stratifying patients by expression quintiles of the corresponding gene. This analysis identified *MYBL2* (alias *B-MYB),* encoding a central transcription factor regulating cell proliferation, cell survival and differentiation^13^, as the top EWSR1-FLI1 upregulated gene, whose high expression in primary EwS was significantly associated with poor OS (nominal *P*=9.6×10^-7^, Bonferroni-adjusted *P*=0.018) (**Fig. 1a,b**).

**Figure 1:**
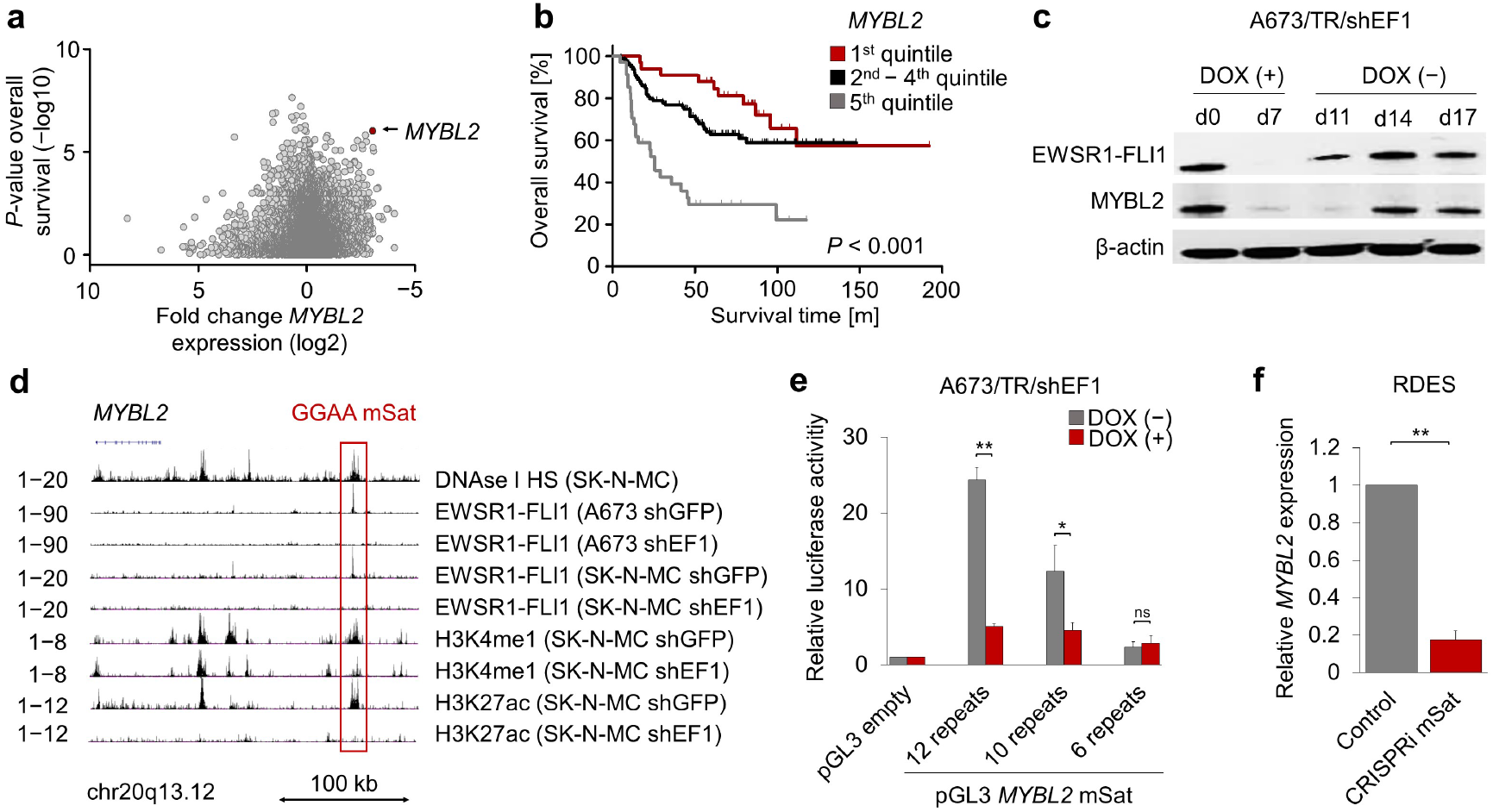
*MYBL2* is a clinically relevant direct EWSR1-FLI1 target gene regulated by a polymorphic GGAA-microsatellite. **a**, Crossing gene expression microarrays of A673 EwS cells, harboring a DOX-inducible shRNA against *EWSR1-FLI1* (A673/TR/shEF1) profiled with/without addition of DOX, with 166 transcriptomes of primary EwS tumors with matched clinical annotation, reveals *MYBL2* as the top EWSR1-FLI regulated gene associated with poor patient outcome. **b**, Kaplan-Meier plot of overall survival in a cohort of 166 primary EwS patients for which Affymetrix gene expression data and clinical annotations were available. Patients were stratified by quintile *MYBL2* expression. Mantel-Haenszel-Test. **c**, Western blot using antibodies against EWSR1-FLI1 and MYBL2 in A673/TR/shEF1 cells. EWSR1-FLI1 was knocked down for 7 days by DOX-treatment and re-expressed by removing DOX from the media for 10 days. β-actin served as loading control. **d**, Epigenetic profile of the *MYBL2* locus inA673 and SK-N-MC EwS cells. Signals from published DNAse-Seq data (DNAse I hypersensitivity; HS) and ChIP-seq data for EWSR1-FLI1, H3K4me1 and H3K27ac in EwS cells transfected with either a control shRNA (shGFP) or a specific shRNA against *EWSR1-FLI1* (shEF1) are displayed. **e**, Luciferase reporter assay after cloning haplotypes of the MYBL2-associated GGAA-microsatellite (mSat) from different cell lines into a pGL3 expression vector. The vectors were transfected in A673/TR/shEF1 cells, which were treated either with or without DOX. Data are mean and SEM, *n=4.* Unpaired student’s t-test. **f**, Analysis of relative *MYBL2* expression in RDES EwS cells after CRISPRi-mediated blockage of protein binding to the MYBL2-associated GGAA-microsatellite. Data are mean and SEM, *n*=4. Unpaired student’s t-test. *** *P*<0.001, ** *P*<0.01, * *P*<0.05.

To validate the EWSR1-FLI1-dependency of MYBL2, we performed time-course experiments in A673/TR/shEF1 revealing that MYBL2 expression followed closely that of EWSR1-FLI1 upon EWSR1-FLI1 knockdown at the mRNA (**Supplementary Fig. 1a**) and protein level (**Fig. 1c**). This EWSR1-FLI1-dependency was confirmed in A673/TR/shEF1 xenografts *in vivo* (**Supplementary Fig. 1b,c**) and 10 additional EwS cell lines *in vitro* (**Supplementary Fig. 1d**).

Despite this tight regulation of MYBL2 by EWSR1-FLI1, we noted a marked inter-tumor heterogeneity of *MYBL2* mRNA expression in 166 primary EwS (**Supplementary Fig. 1e**) and in an independent cohort of 208 primary EwS on protein level stained for p-MYBL2 (**Supplementary Fig. 1f**). Interestingly, *MYBL2* expression did not correlate with minor variations in expression of *EWSR1-FLI1* (**Supplementary Fig. 1g**), suggesting that other factors may cause the inter-individual diversity of *MYBL2* transcription in EwS.

In accordance, re-analysis of published chromatin immunoprecipitation followed by next-generation sequencing (ChIP-seq) data from A673 and SK-N-MC EwS cells revealed strong signals for EWSR1-FLI1 that mapped to a polymorphic GGAA-microsatellite located ~200 kb telomeric of the transcriptional start site of *MYBL2* (**Fig. 1d**). In both cell lines, this GGAA-microsatellite exhibited EWSR1-FLI1-dependent epigenetic characteristics of an active enhancer indicated by H3K4me1 and H3K27ac marks (**Fig. 1d**). The EWSR1-FLI1-dependent enhancer activity of this GGAA-microsatellite was confirmed in reporter assays, in which the enhancer activity of cell line-derived haplotypes comprising 6, 10, and 12 consecutive GGAA-repeats was positively correlated with the number of GGAA-repeats at this GGAA-microsatellite (**Fig. 1e**). Applying the haplotype inference and phasing for short tandem repeats (HipSTR)^14^ algorithm on 18 matched pairs of germline and tumor DNA of EwS patients for which high-quality whole-genome sequencing (WGS) data (150/150bp) covering the MYBL2-associated GGAA-microsatellite were available^8^, we identified additional haplotypes with 6–17 consecutive GGAA-repeats (**Supplementary Table 3**). Of note, all haplotypes (100%, 36/36) were entirely conserved between germline and tumor DNA (**Supplementary Table 3**). Analysis of matched RNA-seq data available for 12 of these 18 EwS primary tumors and one additional tumor sample with WGS and RNA-seq data showed that *MYBL2* expression was higher in tumors with >15 consecutive GGAA-repeats at this microsatellite as compared to tumors with ≤15 GGAA-repeats (**Supplementary Table 3, Supplementary Fig. 1h**).

We further validated the EWSR1-FLI1-mediated regulation of *MYBL2* in time-course EWSR1-FLI1 ChIP-seq and RNA sequencing (RNA-seq) data generated in A673/TR/shEF1 cells. Removal of DOX after long-term suppression of EWSR1-FLI1 for seven days led to a gradual increase of *MYBL2* transcription that correlated with increasing EWSR1-FLI1 recruitment to this GGAA-microsatellite (r_Pearson_=0.816). Strikingly, blockage of protein binding to this *MYBL2-associated* GGAA-microsatellite by CRISPRi (clustered regularly interspaced short palindromic repeats interference), which promotes an inhibiting chromatin state^15,16^, in highly *MYBL2* expressing RDES cells strongly suppressed *MYBL2* transcription (**Fig. 1f**) and induced a potentially counter-regulatory upregulation of *EWSR1-FLI1* (**Supplementary Fig. 1i**). Taken together, these findings indicated that *MYBL2* is a clinically relevant direct EWSR1-FLI1 target gene, whose expression can be modulated by EWSR1-FLI1 binding to an enhancer-like highly polymorphic GGAA-microsatellite.

To obtain first clues on the functional role of MYBL2 in EwS, we performed gene-set enrichment analysis (GSEA) of *MYBL2* co-expressed genes in 166 primary EwS. GSEA revealed that *MYBL2* co-expressed genes were significantly enriched in human orthologs of known MYBL2 targets in zebrafish^17^ as well as in signatures related to proliferation^18^, cell cycle progression^19^, and sensitization to apoptosis mediated by a CDK-inhibiting protein^20^ (min. nominal enrichment score (NES)=2.76, *P*<0.001) (**Fig. 2a, Supplementary Table 4**), suggesting that MYBL2 may constitute a key downstream mediator of EWSR1-FLI1-induced, evolutionary conserved proliferation programs.

**Figure 2:**
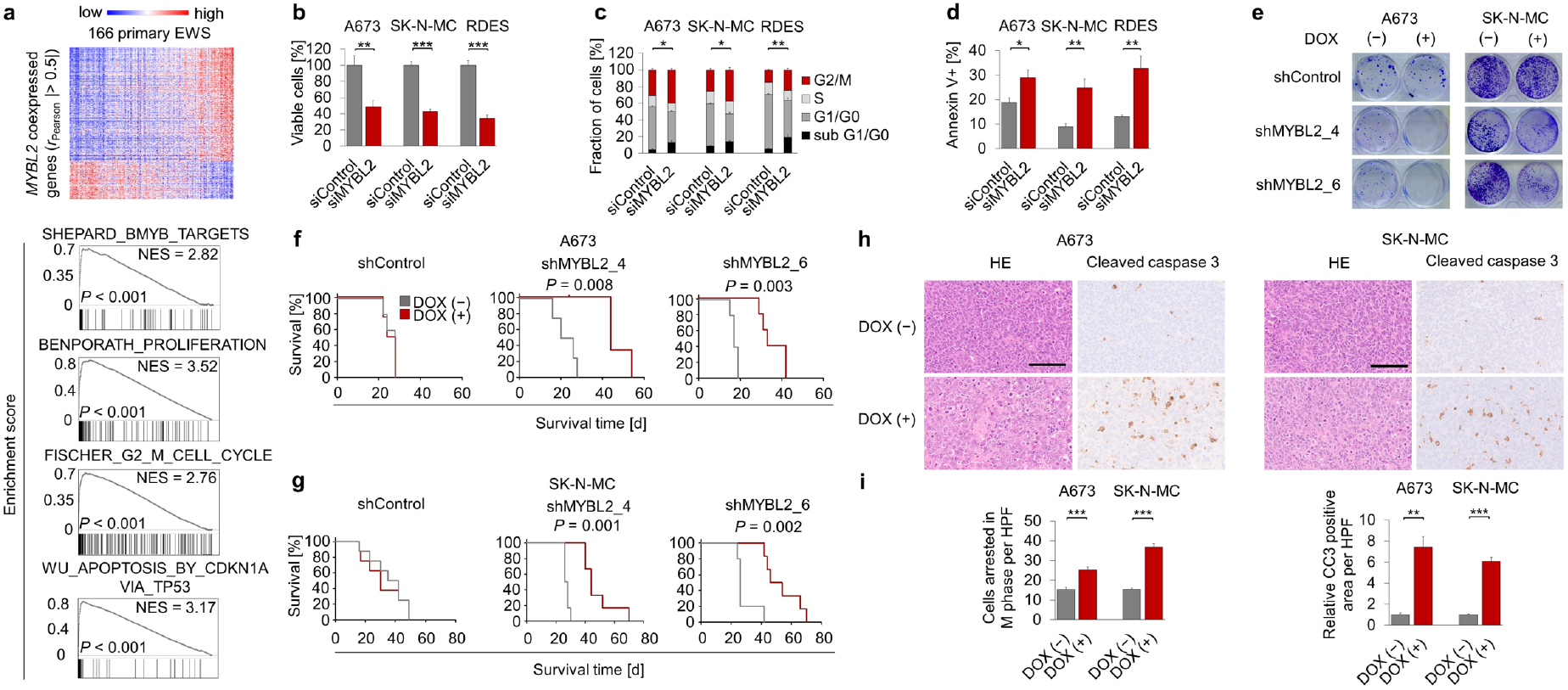
MYBL2 is critical for proliferation and cell survival of EwS cells *in vitro* and *in vivo.* **a**, Heat-map showing genes whose expression is positively or negatively correlated with *MYBL2* expression in Affymetrix gene expression data of 166 primary EwS tumors. GSEA of the same dataset showing selected gene sets enriched in *MYBL2* high-expressing tumors, which comprised MYBL2 targets^17^, proliferation^18^, cell cycle progression^19^ and sensitization to apoptosis mediated by a CDK-inhibiting protein^20^. **b**, Viable cell count 96h after transfection of three different EwS cell lines with either four different specific siRNAs directed against *MYBL2* (summary of four different siRNAs shown) or a non-targeting siControl. Data are mean and SEM, *n*≥3. Unpaired student’s t-test. **c**, Analysis of cell cycle phases and cell death (sub G1/G0) 96h after transfection with either four different specific siRNAs directed against *MYBL2* (summary of four different siRNAs shown) or a nontargeting siControl, using Propidium-Iodide staining and flow cytometry. Data are mean and SEM, n=3. Unpaired student’s t-test. **d**, Analysis of apoptosis in three different EwS cell lines 96h after transfection with either four different specific siRNAs directed against *MYBL2* (summary of four different siRNAs shown) or a non-targeting siControl, using annexin V/propidium-iodide staining and flow cytometry. Data are mean and SEM, n=3. Unpaired student’s t-test. **e**, Representative colony forming assays (CFAs) of A673 and SK-N-MC cells containing either DOX-inducible specific shRNA constructs directed against *MYBL2* or a non-targeting shControl. Cells were grown either with or without DOX. Colonies were stained with Crystal-violet 10–14 days after seeding. **f,g**, Kaplan-Meier plots showing survival time of NSG mice xenografted with A673 or SK-N-MC cells, containing either DOX-inducible specific shMYBL2 or non-targeting shControl constructs. Once tumors were palpable, mice were randomized and treated with either vehicle (−) or DOX (+). Mantel-Haenszel-Test. **h**, Representative micrographs of HE and IHC staining for cleaved caspase 3 (CC3) of NSG mice xenografted with A673 or SK-N-MC cells containing a DOX-inducible specific shMYBL2 Mice were treated with either vehicle (−) or DOX (+). Scale bar is 50 μm. **i**, Quantification of cells arrested in M-phase and automated quantification of the picture area positive for CC3 of xenografts from A673 or SK-N-MC cells containing a DOX-inducible specific shMYBL2. Mice were treated with either vehicle (−) or DOX (+). Data are mean and SEM of ten high-power fields in five tumors for each condition. Unpaired student’s t-test. *** *P*<0.001, ** *P*<0.01, * *P*<0.05.

To test this hypothesis, we performed siRNA-mediated *MYBL2* knockdown experiments in three different EwS cell lines (A673, SK-N-MC, and RDES) with moderate to high baseline *MYBL2* expression (**Supplementary Fig. 2a**). Using four different siRNAs (**Supplementary Fig. 2b, c**), we found that knockdown of *MYBL2* significantly reduced cell proliferation through blockage of G2/M-progression, which was accompanied by apoptotic cell death (**Fig. 2b-2d**). To further explore the function of MYBL2 in EwS growth, we generated two EwS cell lines (A673 and SK-N-MC) with DOX-inducible anti-MFBL2 shRNA expression systems using two different shRNAs (**Supplementary Fig. 2d**). In both cell lines, DOX-induced *MYBL2* silencing significantly reduced clonogenic growth *in vitro* (**Fig. 2e, Supplementary Fig. 2e**) and tumor growth *in vivo* (**Fig. 2f,g, Supplementary Fig. 2f,g**) compared to a nontargeting control shRNA. Consistent with our transient knockdown experiments (**Fig. 2b-d**), we observed an increased number of stalled mitoses, indicating G2/M-blockage, and more apoptotic tumor cells positive for cleaved caspase 3 in xenografts with *MYBL2* suppression (**Fig. 2h,i**). Collectively, these findings indicate that MYBL2 is a critical pro-proliferative downstream effector of EWSR1-FLI1 required for proper G2/M-transition and cell survival.

To identify potential direct MYBL2 targets that could explain its strong pro-proliferative effect, we sequenced RNA of three EwS cell lines with/without siRNA-mediated knockdown of *MYBL2* (**Fig. 3a**). Consistent with our enrichment analyses in primary EwS and functional experiments, GSEA of the identified differentially expressed genes (DEGs) showed that *MYBL2* suppression was accompanied by a significant downregulation of the same gene-set comprising human orthologs of zebrafish MYBL2 targets and identical proliferation, cell cycle and sensitization to CDK-inhibitor mediated apoptosis gene signatures (max. NES=-1.89, *P*<0.001) (**Fig. 3b, Supplementary Fig. 3a, Supplementary Table 5**). We then focused on the 76 most significantly DEGs (mean log2 FC |≥1.5|, Bonferroni-adjusted *P*<0.05) (**Supplementary Fig. 3b, Supplementary Table 6**), of which representative genes were validated by qRT-PCR (**Supplementary Fig. 3c**). Subsequent ChIP-seq analysis using a specific anti-MYBL2 antibody proved that 66% of these DEGs (50/76) exhibited strong MYBL2 binding at their promoters (**Fig. 3c, Supplementary Table 7**). Using microarray data of 166 primary EwS, generated with platforms on which 92% (46/50) direct MYBL2 target genes were represented, we correlated the expression levels of these genes with that of *MYBL2* (**Supplementary Table 8**) and these genes’ potential association with patients’ OS stratifying patients by median expression of the corresponding gene (**Supplementary Table 9**). Among these genes, *CCNF, BIRC5* and *AURKB* stood out for being highly significantly co-expressed with *MYBL2* (Bonferroni-adjusted *P*<0.05, r_Pearson_≥0.7) (**Fig. 3d**), and associated with poor OS (P<0.05) (**Fig. 3e**). To investigate their functional role, we individually knocked down either gene using two specific siRNAs in two different EwS cell lines (**Supplementary Fig. 3d**) and assessed proliferation and cell viability *in vitro*. Strikingly, knockdown of these genes broadly phenocopied the anti-proliferative and antisurvival effect of *MYBL2* silencing (**Fig. 3f,g**), suggesting that they may constitute important mediators of the pro-proliferative EWSR1-FLI1/MYBL2 transcriptional program.

**Figure 3:**
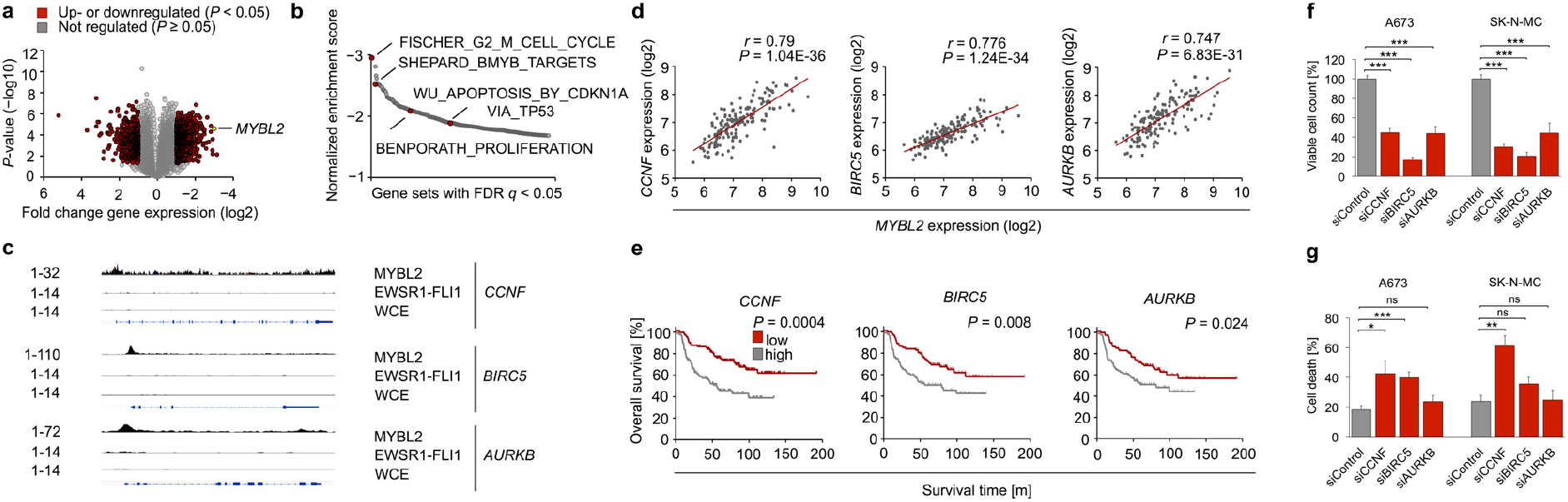
MYBL2 mediates its phenotype via direct upregulation of *CCNF, BIRC5* and *AURKB.* a, Analysis of RNA-seq data showing differentially expressed genes (DEGs) after siRNA-mediated *MYBL2* knockdown compared to a non-targeting siControl. A summary of three cell lines is shown; *n*=3 biological replicates per group per cell line. **b**, GSEA of RNA-seq data shown in (a). Displayed are 275 downregulated gene sets that had an FDR q<0.05. **c**, Analysis of MYBL2 ChIP-seq data from A673 cells shows MYBL2 peaks in the promoter regions of *CCNF, BIRC5* and *AURKB.* Publicly available EWSR1-FLI1 ChIP-seq data from A673 cells was analyzed to exclude a direct regulation by EWSR1-FLI1 of these three genes. Whole cell extract (WCE) served as a control. **d**, Linear regression of *CCNF*, *BIRC5* or *AURKB* expression onto *MYBL2* expression in an Affymetrix gene expression dataset of 166 primary EwS tumors. **e**, Analysis of overall patient survival in a cohort of 166 primary EwS patients for which Affymetrix gene expression data and clinical annotations were available. Patients were stratified by median expression levels of the indicated gene. Mantel-Haenszel-Test. **f**, Viable cell count 96h after transfection of A673 and SK-N-MC cell lines with two different siRNA directed against either *CCNF, BIRC5, AURKB* (summary of two different siRNAs shown) or a non-targeting siControl. Data are mean and SEM, n=3. Unpaired student’s t-test. **g**, Cell death as measured by trypan blue positivity 96h after transfection of A673 and SK-N-MC EwS cells with two different siRNA directed against either *CCNF, BIRC5, AURKB* (summary of two different siRNAs shown) or a non-targeting siControl. Data are mean and SEM, *n*=3. Unpaired student’s t-test. *** *P*<0.001, ** *P*<0.01, * *P*<0.05.

As there are – to the best of our knowledge – currently no direct MYBL2 inhibitors available, we reasoned that targeting its major upstream cyclin dependent kinase, CDK2, which activates MYBL2 through phosphorylation^13^, may offer a new therapeutic option for EwS patients with high MYBL2 expression. To test this possibility, we treated EwS cells with two different low molecular weight CDK2 inhibitors (CVT-313 and NU6140). While both inhibitors showed strongly reduced growth of A673 EwS cells at the lower micro-molar range (**Fig. 4a**), the sensitivity toward both inhibitors was dramatically diminished when *MYBL2* was suppressed (**Fig. 4a**). Such differential effect was not observed in control cells expressing a non-targeting shRNA (**Fig. 4a**). Treatment of NOD/scid/gamma (NSG) mice with NU6140 significantly reduced growth of EwS xenografts as compared to vehicle (DMSO) (**Fig. 4b**), and was accompanied by reduced levels of phosphorylated MYBL2 as evidenced by immunohistochemistry (**Fig. 4c**). However, CDK2-inhibition showed no additional effect on growth of xenografts with silenced *MYBL2* expression (**Fig. 4b**), suggesting that MYBL2 is required for the anti-proliferative effect of CDK2 inhibitors. Consistently, different EwS cell lines with high *MYBL2* levels showed higher sensitivity to treatment with NU6140 as compared to a EwS cell line with constitutively low *MYBL2* levels (**Supplementary Fig. 4a,b**). Since we neither observed significant weight loss (**Supplementary Fig. 4c**) nor histo-morphological changes in inner organs in mice treated for 16 days with up to 40 mg/kg, these results indicated that CDK2-inhibition can safely impair growth of EwS tumors and that MYBL2 may serve as a biomarker to predict their efficacy.

**Figure 4:**
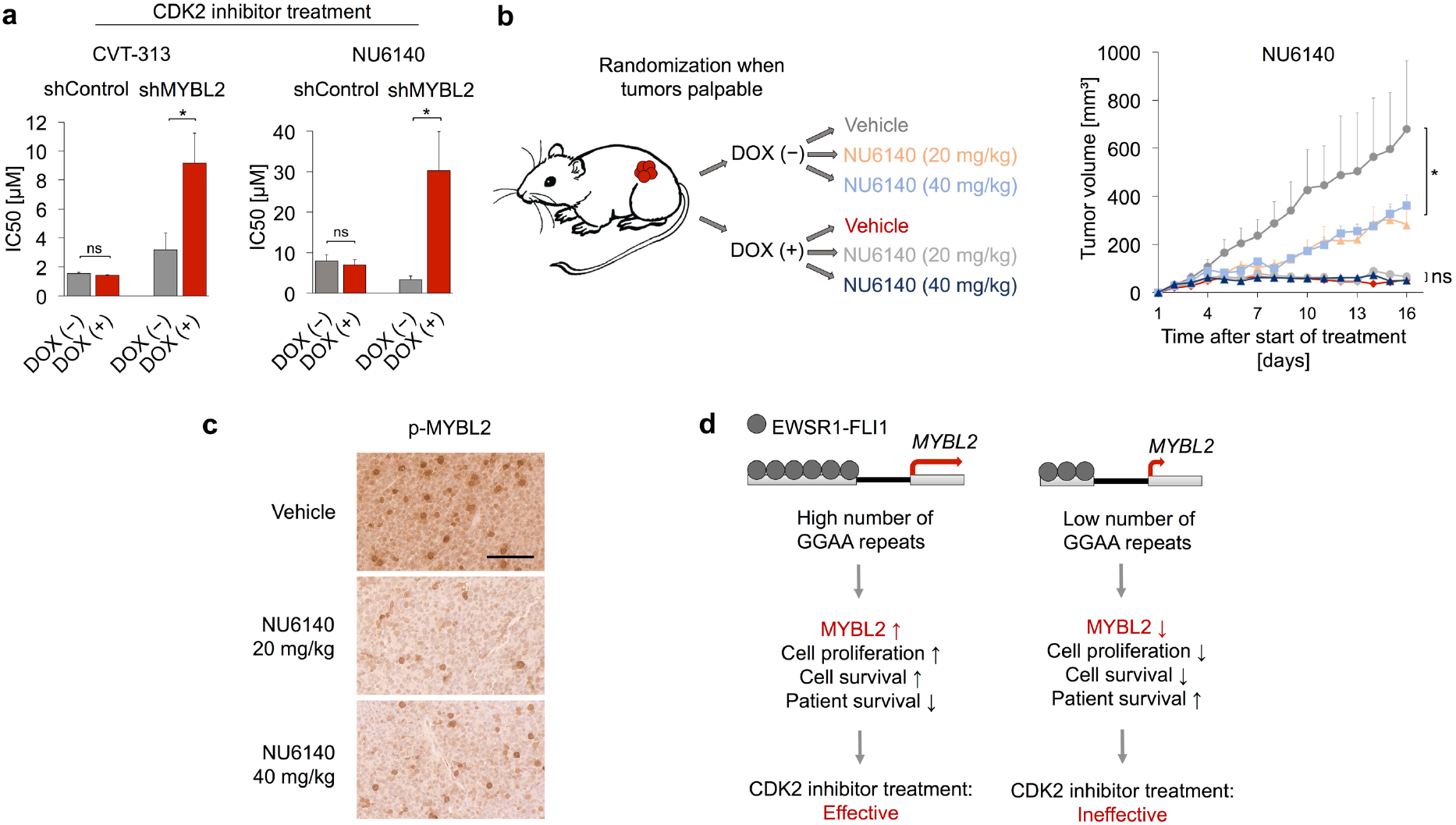
High MYBL2 expression levels sensitize EwS cells toward treatment with CDK2 inhibitors *in vitro* and *in vivo.* a, A673 cells containing either DOX-inducible specific shMYBL2 or non-targeting shControl constructs were treated with the CDK2 inhibitors CVT-313 or NU6140 at different concentrations. Additionally, cells were treated either with or without DOX. Relative cell viability was assessed by a Resazurin assay. Data are mean and SEM, n≥3. Unpaired student’s t-test. **b**, Left: Schematic of the experimental setting of CDK2 inhibitor treatment (NU6140) *in vivo* and tumor volume curves of NSG mice xenografted with A673 cells containing a DOX-inducible shMYBL2 construct, treated with/without DOX and either vehicle or NU6140 in a dose of 20 mg/kg or 40 mg/kg. Right: Mice were randomized to the treatment groups as soon as tumors were palpable. For each condition the mean tumor volume and SEM of 3–6 mice is shown. Unpaired student’s t-test. **c**, Representative micrographs of IHC staining for p-MYBL2 of NSG mice xenografted with A673/TR/shMYBL2 cells without DOX, but treatment with either vehicle or CDK2 inhibitor at a dose of 20 mg/kg or 40 mg/kg. Scale bar is 50 μm. **d**, Model of EWSR1-FLI1-dependent regulation of *MYBL2* in EwS and its functional and therapeutic implications. *** *P*<0.001, ** *P*<0.01, * *P*<0.05.

Collectively, our results demonstrate that oncogenic cooperation between a dominant oncogene (here EWSR1-FLI1) and a regulatory germline variant (here a polymorphic enhancer-like GGAA-microsatellite) could be a major source of inter-tumor heterogeneity determining outcome and drug response of an aggressive childhood cancer through modulation of a druggable key downstream player (**Fig. 4e**). These results also suggest that cooperation between disease-promoting somatic mutations and regulatory germline variants could constitute a general mechanism to explain diversity of disease phenotypes, possibly beyond cancer. In line with this idea, recent reports for neurodegenerative and metabolic diseases showed that the same disease-causing somatic event/mutation can lead to distinct phenotypes depending on (inherited) variations in the regulatory genome^6,21,22^. We anticipate that our findings made in the EwS model are translatable to other malignancies such as ETS-driven leukemia and TMPRSS2-ETS-driven prostate carcinoma^23^, and propose that integration of the regulatory genome in the process of developing new diagnostic and/or therapeutic strategies is necessary to refine and fully exploit ‘omics’-based precision medicine.

## MATERIALS AND METHODS

### Provenience of cell lines and cell culture conditions

A673 and HEK293T cells were purchased from American Type Culture Collection (ATCC). RDES, SK-N-MC, and MHH-ES1 cells were provided by the German collection of Microorganisms and Cell lines (DSMZ). TC-32, TC-71, CHLA-10 cells were kindly provided by the Children’s Oncology Group (COG) and EW1, EW7, EW16, EW18, EW22, ES7, ORS, POE, STA-ET1 cells were provided by O. Delattre (Institute Curie, Paris). A673/TR/shEF1 cells were kindly provided by J. Alonso (Madrid, Spain)^24^. All cell lines were grown at 37°C and 5% CO2 in humidified atmosphere. RPMI 1640 medium supplemented with stable glutamine (Biochrom), 10% tetracycline-free FCS (Sigma-Aldrich), 100 U/ml penicillin (Biochrom), and 100 μg/ml streptomycin (Biochrom) was used to grow the cells. Cells were routinely checked by nested PCR for mycoplasma infection. Cell line purity was confirmed by STR-profiling.

### DNA/RNA extraction, reverse transcription and quantitative real-time PCR (qRT-PCR)

DNA was extracted with the NucleoSpin Tissue kit (Macherey-Nagel), plasmid DNA was extracted from bacteria with the PureYield kit (Promega). RNA was extracted with the NucleoSpin II kit (Macherey-Nagel) and reverse-transcribed using the High-Capacity cDNA Reverse Transcription kit (Applied Biosystems). Genomic qRT-PCRs were performed using SYBR green (Applied Biosystems). Oligonucleotides were purchased from MWG Eurofins Genomics. Reactions were run on a Bio-Rad CFX Connect instrument and analyzed using the Bio-Rad CFX Manager 3.1 software. For primer sequences see **Supplementary Table 10**.

### Transient transfection

For siRNA transfection, cells were seeded in a six-well plate at a density of 1.5×10^5^ per well in 1.6 ml of growth medium. The cells were transfected with either a negative control non targeting siRNA (Sigma-Aldrich MISSION siRNA Universal Negative Control #1) or specific siRNAs (25 to 65 nM, depending on the cell line and the siRNA) and HiPerfect (Qiagen). Cells were retransfected 48h after the first transfection and harvested 96h after the first transfection. siRNA sequences are given in **Supplementary Table 10**.

For plasmid transfection, cells were seeded in a six-well plate at a density of 2× 10^5^ per well in 1.8 ml of growth medium. Plasmids were transfected with Lipofectamine LTX and Plus Reagent (Invitrogen). The pGL3 vector used for reporter assays has been described before^9^.

### Doxycycline (DOX)-inducible shRNA constructs

Either a non-targeting negative control shRNA (MWG Eurofins Genomics) or specific shRNAs targeting *MYBL2* (MWG Eurofins Genomics) were cloned in the pLKO-Tet-on-all-in-one system^25^. Oligonucleotide sequences are given in **Supplementary Table 10**. Lentivirus production was performed in HEK293T cells. A673 and SK-N-MC EwS cells were infected with the respective lentiviruses and selected with 1.5 μg/ml puromycin (Invivogen). After single cell cloning, knockdown efficacy of individual clones was assessed by qRT-PCR 48h after addition of DOX (1 μg/ml; Sigma-Aldrich).

### DNA constructs and reporter assays

MYBL2-associated microsatellites (with 500 bp 5′ and 3′ flanking regions) from three EwS cell lines were PCR-cloned upstream of the SV40 minimal promoter into the pGL3-luc vector (Promega)^9^. Primer sequences are given in **Supplementary Table 10**. A673/TR/shEF1 cells (2×10^5^ per well) were transfected with the microsatellite-containing pGL3-luc vectors and *Renilla* pGL3-Rluc vectors (ratio, 100:1) in a six-well plate with 1.8 ml of growth medium. Transfection media was replaced by media with/without DOX (1 μg/ml) 4h after transfection. After 72h the cells were lysed and assayed with a dual luciferase assay system (Berthold). Firefly luciferase activity was normalized to Renilla luciferase activity.

### Generation of cell lines with inducible CRISPR interference (CRISPRi)

Due to the lack of functional DNAse, CRISPRi does not cause a knockout of the targeted DNA sequence, but blocks protein binding to it^15,16^. For the reported experiments, a DNAse-dead CAS9 (dCAS9) fused to the KRAB effector domain, which promotes an inhibiting chromatin state, is targeted to the genomic region of interest by specific gRNAs to silence the activity of a given enhancer^15,16^. To achieve this, we used a pHAGE TRE dCas9-KRAB vector (Addgene #50917) and a pLKO.1-puro U6 sgRNA BfuAI large stuffer vector (Addgene #52628), the latter one containing either two gRNAs, targeting sequences adjacent to the MYBL2-associated GGAA-microsatellite or a scrambled control (**Supplementary Table 10**). Lentivirus production was performed in HEK293T cells. RDES EwS cells were infected with the respective lentiviruses and selected with 1 μg/ml puromycin (Invivogen) and 1. 5 μg/ml G418 (Invivogen). The cells were induced with DOX for five days, after which *MYBL2* and *EWSR1-FLI1* levels were measured by qRT-PCR.

### Western blot

Protein from A673/TR/shEF1 cells was extracted at d0, d7, d11, d14 and d17 with RIPA and anti-protease cocktail (Roche). Western blots were performed following routine protocols and specific band detection was achieved by the use of rabbit monoclonal anti-FLI1 antibody (1:1,000, ab133485, Abcam), rabbit polyclonal anti-MYBL2 antibody (1:500, sc-725, Santa Cruz) and mouse monoclonal anti-β-actin (1:10,000, A-5316, Sigma-Aldrich). Anti-rabbit IgG horseradish peroxidase coupled antibody (1:3,000, Amersham Bioscience) and antimouse IgG horseradish peroxidase coupled antibody (1:3,000, Amersham Bioscience) was used as secondary antibody. Proteins were visualized using chemiluminescence (Pierce ECL Western blot chemiluminescent substrate (Thermo Fisher Scientific).

### Proliferation assays

Cells were seeded in a six-well plate at a density of 1.5×10^5^ per well in 1.6 ml of growth medium. The cells were transfected with either a negative control non-targeting siRNA (Sigma-Aldrich MISSION siRNA Universal Negative Control #1) or specific siRNAs (25 to 65 nM, depending on the cell line and the siRNA) using HiPerfect (Qiagen). siRNA sequences are given in **Supplementary Table 10**. Retransfection was performed 48h after the first transfection. 96h after the first transfection, cells were harvested (including supernatant), stained with Trypan blue (Sigma-Aldrich) and counted in a standardized hemocytometer (C-chip, NanoEnTek).

### Analysis of cell cycle and apoptosis

Analysis of cell cycle phases was performed by propidium iodide (PI) (Sigma-Aldrich) staining. Cells were transfected with siRNAs equivalently to the proliferation assays (see above), harvested after 96h (including supernatant), fixed in ethanol (70%) at 4°C, and stained with PI solution (50 μg/ml, with 20 μg/ml RNAse A). Analysis of apoptosis has been performed by combined Annexin V-FITC/PI staining (BD Pharmingen FITC Annexin V Apoptosis Detection Kit II). Cells were transfected with siRNAs equivalently to the proliferation assays (see above) and harvested after 96h (including supernatant). The samples were assayed on an Accuri C6 flow cytometer and analyzed with the Accuri C6 CFlow Plus software.

### Clonogenic growth assays (Colony forming assays)

A673 and SK-N-MC cells containing either a doxycycline-inducible non-targeting control shRNA or MYBL2-targeting specific shRNAs were seeded in a 12-well plate at a density of 500 cells (A673) or 1000 cells (SK-N-MC) per well in 2 ml of growth medium. Cells were grown with/without DOX (1 μg/ml) for 10-14 days depending on the cell line and afterwards stained with crystal violet (Sigma-Aldrich). Colony number was determined on scanned plates with Fiji (ImageJ)^26,27^.

### cDNA library and RNA sequencing (RNA-seq)

A673, SK-N-MC and RDES EwS cell lines were transfected with either a negative control non-targeting siRNA or a specific siRNAs targeting *MYBL2.* Total RNA was extracted using the NucleoSpin II kit (Macherey-Nagel). Complementary DNA libraries were sequenced with an Illumina HiSeq2500 instrument using 150bp paired-end sequencing. Obtained reads were aligned on the human genome (hg19) using TopHat (version 2.0.6)^28^. Counting of reads on annotated genes from the GRCh37 gene build was done using htseq-count (v. HTSeq-0. 5.3p9)^29^ with the following parameters: htseq-count -a 10 -q -s no -m union. Sample-to-sample normalization and differential expression analyses were performed using the R package DESeq2 (v. 1.18.0)^30^. RNA-seq data were deposited at the Gene Expression Omnibus (GEO; accession code GSE119972).

### Chromatin immunoprecipitation and sequencing (ChIP-seq)

DNA-protein cross-linking was done in presence of 1% of paraformaldehyde on 12×10^6^ A673 cells for each condition for 10 min. Cell lysis, chromatin shearing, immunoprecipitation and DNA purification was performed with reagents from iDeal ChIP-seq kit for Transcription Factors (Diagenode, ref: C01010054). Chromatin shearing was carried out in a Bioruptor (Diagenode) using 20 cycles of sonication (30 sec high, 30 sec off) in TPX tubes (Diagenode, ref: 50001). For immunoprecipitation of MYBL2, 2 μg of a monoclonal ChIP-grade rabbit anti-MYBL2 antibody (Abcam, ref: 76009) were used. IgG and CTCF ChIP was included as negative and positive control, respectively. MYBL2 ChIP and input were sequenced on a HiSeq2500 instrument (Illumina) using 100bp single-end sequencing. ChIP-seq reads were aligned to the human genome (hg19 version) with Bowtie2^31^. Peaks were called with MACS2 with option narrow^32^. To normalize, we took the input dataset from the same cell line. PAVIS was used for peak annotation and visualization^33^. ChIP-seq data were deposited at the GEO (accession code GSE119972).

### Analysis of published ChIP-seq and DNase-seq data

Publicly available ENCODE SK-N-MC DNase-seq data (GSM736570)^34^ and pre-processed A673 and SK-N-MC ChIP-seq data (GSE61944)^35^ were retrieved from the GEO and displayed in the UCSC genome browser. Samples used: GSM1517544 SK-N-MC_shGFP_48h_FLI 1; GSM1517553 SK-N-MC_shFLI1_48h_FLI1; GSM1517569 A673_shGFP_48h_FLI1; GSM1517572 A673_shFLI1_48h_FLI1; GSM1517548 SK-N-MC_shGFP_96h_H3K4me 1; GSM1517557 SK-N-MC_shFLI1_96h_H3K4me1; GSM1517545 SK-N-MC_shGFP_48h_H3K27ac; GSM1517554 SK-N-MC_shFLI1_48h_H3K27ac; GSM1517568 A673 whole cell extract (WCE).

### CDK2 inhibitor assays *in vitro*

Cells were seeded in a 96-well plate at a density of 5×10^3^ per well with/without DOX (1 μg/ml). After 24h of pre-incubation with DOX, CDK2 inhibitors (CVT-313 or NU6140 (Merck)) were added in serially diluted concentrations ranging from 100 nM to 0.001 nM. Every well contained an equal concentration of 0.5% DMSO. Cells only treated with 0.5% of DMSO served as a control. After 72h of inhibitor treatment, the plates were assayed on a Thermo Fisher Varioskan plate reader after adding Resazurin (20 μg/ml) for 6h.

### Xenotransplantation experiments and CDK2 inhibitor treatment *in vivo*

3×10^6^ A673 and SK-N-MC cells, containing either a DOX-inducible negative control shRNA or specific shRNAs against *MYBL2,* were injected subcutaneously with a 1:1 mix of PBS (Biochrom) and Geltrex (LDEV-Free Reduced Growth Factor Basement Membrane Matrix,

Thermo Fisher Scientific; max volume 100 μl) in the right flanks of 3-9 months old NSG mice (Charles River Laboratories). For shRNA sequences see **Supplementary Table 10**. When tumors were first palpable, mice were randomized to the control group (17.5 mg/ml sucrose (Sigma-Aldrich) in drinking water) or the treatment group (2 mg/ml DOX (Beladox, bela-pharm) and 50 mg/ml sucrose (Sigma-Aldrich) in drinking water). Tumor size was measured with a caliper every two days and tumor volume was calculated as *V=a×b^2^/2* with *a* being the largest diameter and *b* the smallest. Once the tumors reached a volume of 1,500 mm^3^ respective mice were sacrificed by cervical dislocation. For CDK2 inhibitor treatment *in vivo,* cells were injected as described above. When tumors were palpable, mice were assigned to either the vehicle or treatment group (20 mg/kg and 40 mg/kg), each with or without addition of DOX to the drinking water (2 mg/ml DOX; Beladox, bela-pharm). The CDK2 inhibitor (NU6140, Bio-Techne) was administered i.p. for 12 days, with a break of one day every 4 days of treatment. To check histomorphological changes of inner organs upon CDK2 inhibitor treatment we examined hematoxylin and eosin (HE) stained slides of heart, lungs, liver, stomach, pancreas, intestines, kidneys, adrenal glands, bone marrow and spleen from treated vs. non-treated mice. Experiments were approved by local authorities and conducted in accordance with the recommendations of the European Community (86/609/EEC) and UKCCCR (guidelines for the welfare and use of animals in cancer research). The sample size was not predetermined.

### Survival analysis

Microarray data of 166 primary EwS tumors (GSE63157^36^, GSE34620^37^, GSE12 1 02^38^, GSE17618^39^) for which well-curated clinical annotations were available were downloaded from the GEO. The data were either generated on Affymetrix HG-U133Plus2.0 or on Affymetrix HuEx-1.0-st microarray chips and were normalized separately by RMA using custom brainarray chip description files (CDF, v20). Batch effects were removed using

ComBat^40,41^. Samples were stratified by their quintile or median intra-tumoral gene expression levels. Significance levels were calculated with a Mantel-Haenszel test (calculated using GraphPad Prism version 5). *P* values <0.05 were considered as statistically significant. Survival data were crossed with gene expression microarray data (Affymetrix HG-U133A2.0) generated in A673/TR/shEF1 cells (GSE27524; 53h DOX-treatment)^42^, which were normalized as described above (brainarray CDF, v19).

### Gene-set enrichment analysis (GSEA)

Using the Affymetrix gene expression dataset comprising 166 primary EwS patients, enrichment of gene sets that are among *MYBL2* co-regulated genes were identified by ranking of Pearson’s correlation coefficient of the expression of every gene with *MYBL2* expression and performance of a pre-ranked GSEA with 1,000 permutations^43^. Using the RNA-seq dataset containing DEGs after siRNA-mediated *MYBL2* knockdown compared to a nontargeting siControl in A673, SK-N-MC and RDES EwS cell lines, all genes were ranked by their mean log2 FC and a pre-ranked GSEA was performed with 1,000 permutations^43^.

### GGAA-microsatellite analysis of matched germline and tumor DNA pairs using HipSTR

EwS tumors and/or matched blood samples were collected from 19 EwS patients treated in the Hospital for Sick Children (SickKids) in Toronto, Canada, in accordance with Research Ethical Board (REB) guidelines. Whole-genome sequencing was performed in all tumors and in available matched 18 blood samples using established protocols on Illumina instruments (paired end 150/150bp) with PCR-free specifications for four of 19 samples. Paired-end FASTQ files were aligned to the human genome (hg19/GRCh37) using BWA-MEM (v.0.7.8). Indel realignment and base quality scores were recalibrated using the Genome Analysis Toolkit (v.2.8.1). In addition, RNA-seq was carried out in 13 of the 19 EwS tumors, as described^8^. To call the genotypes of the MYBL2-associated GGAA-microsatellite, we applied HipSTR (v.0.6.2)^14^ on the 18 tumor/normal pairs^8^ using a minimum threshold of ten reads. All genotypes passed the following HipSTR default filters: --min-call-qual 0.9; --max-call-flank-indel 0.15; --max-call-stutter 0.15; --min-call-allele-bias −2; --min-call-strand-bias – 2.

### Human samples and ethics approval

Human tissue samples were retrieved from the archives of the Institute of Pathology of the LMU Munich (Germany) and the Gerhard-Domagk Institute of Pathology of the University Hospital of Münster (Germany) with approval of the corresponding institutional review boards.

### Construction of tissue microarrays (TMA) and immunohistochemistry (IHC)

The composition and construction of the used TMAs was described previously^44^. All EwS FFPE samples showed cytogenetic evidence for a translocation of the *EWSR1* gene as determined by fluorescence in situ hybridization (FISH) and/or qRT-PCR and were reviewed by a reference pathologist. Each TMA slide contained at least 2 cores, each 1 mm in diameter, from every sample as well as internal controls. For IHC, 4 μm sections were cut and antigen retrieval was performed with microwave treatment using the antigen retrieval ProTaqs I Antigen-Enhancer (Quartett) for p-MYBL2 or the Target Retrieval Solution (Agilent Technologies) for cleaved caspase 3. Blockage of endogenous peroxidase was performed using 7.5% aqueous H2O2 solution at room temperature and blocking serum from the corresponding kits for 20 min. Slides were then incubated for 60 min with the primary antibodies anti-p-MYBL2 (1:100 dilution, Abcam, ab76009) and cleaved caspase 3 (1:100 dilution, Cell signaling, #9661). Then slides were incubated with a secondary anti-rabbit IgG antibody (MP-7401, ImmPress Reagent Kit, Peroxidase-conjugated) followed by target detection using DAB+ chromogen (Agilent Technologies). Slides were counterstained with hematoxylin Gill’s Formula (H-3401, Vector).

### Evaluation of immunoreactivity

Semi-quantitative evaluation of marker immunostaining was carried out by an independent observer in analogy to scoring of hormone receptor Immune Reactive Score (IRS) ranging from 0-12 as described^45^. The percentage of cells with high nuclear expression was scored in five tiers (score 0=0%, score 1=0–9%, score 2=10–50%, score 3=51–80% and score 4=81–100%). The fraction of cells showing high nuclear expression was scored after examination of 10 high-power fields (40×) of at least one section for each sample. In addition, the intensity of marker immunoreactivity was determined (score 0=none, score 1=low, score 2=intermediate and score 3=strong). The product of both scores defined the final IRS.

## FUNDING

The laboratory of T. G. P. G. is supported by grants from the ‘Verein zur Förderung von Wissenschaft und Forschung an der Medizinischen Fakultät der LMU München (WiFoMed)’, by LMU Munich’s Institutional Strategy LMUexcellent within the framework of the German Excellence Initiative, the ‘Mehr LEBEN für krebskranke Kinder – Bettina-Bräu-Stiftung’, the Walter Schulz Foundation, the Wilhelm Sander-Foundation (2016.167.1), the Friedrich-Baur foundation, the Matthias-Lackas foundation, the Dr. Leopold und Carmen Ellinger foundation, the Gert & Susanna Mayer foundation, the Deutsche Forschungsgemeinschaft (DFG 391665916), and by the German Cancer Aid (DKH-111886 and DKH-70112257). J. L. was supported by a scholarship of the China Scholarship Council (CSC). J. M. was supported by a scholarship of the ‘Kind-Philipp-Foundation’, M. D. by a scholarship of the ‘Deutsche Stiftung für Junge Erwachsene mit Krebs’, and T.L.B.H. by a scholarship of the German Cancer Aid. M. F. O. and M. M. L. K. were supported by scholarships of the German National Academic Foundation. M. C. B. was supported by a scholarship of the Max Weber-Program of the State of Bavaria.

## ACKNOWLEGEMENTS

We thank Dr. Sarah-Maria Fendt for critical reading of our manuscript, and Anja Heier and Andrea Sendelhofert for excellent technical assistance.

## CONFLICT OF INTEREST

We declare no conflicts of interest.

---

**Supplementary Figure 1:**
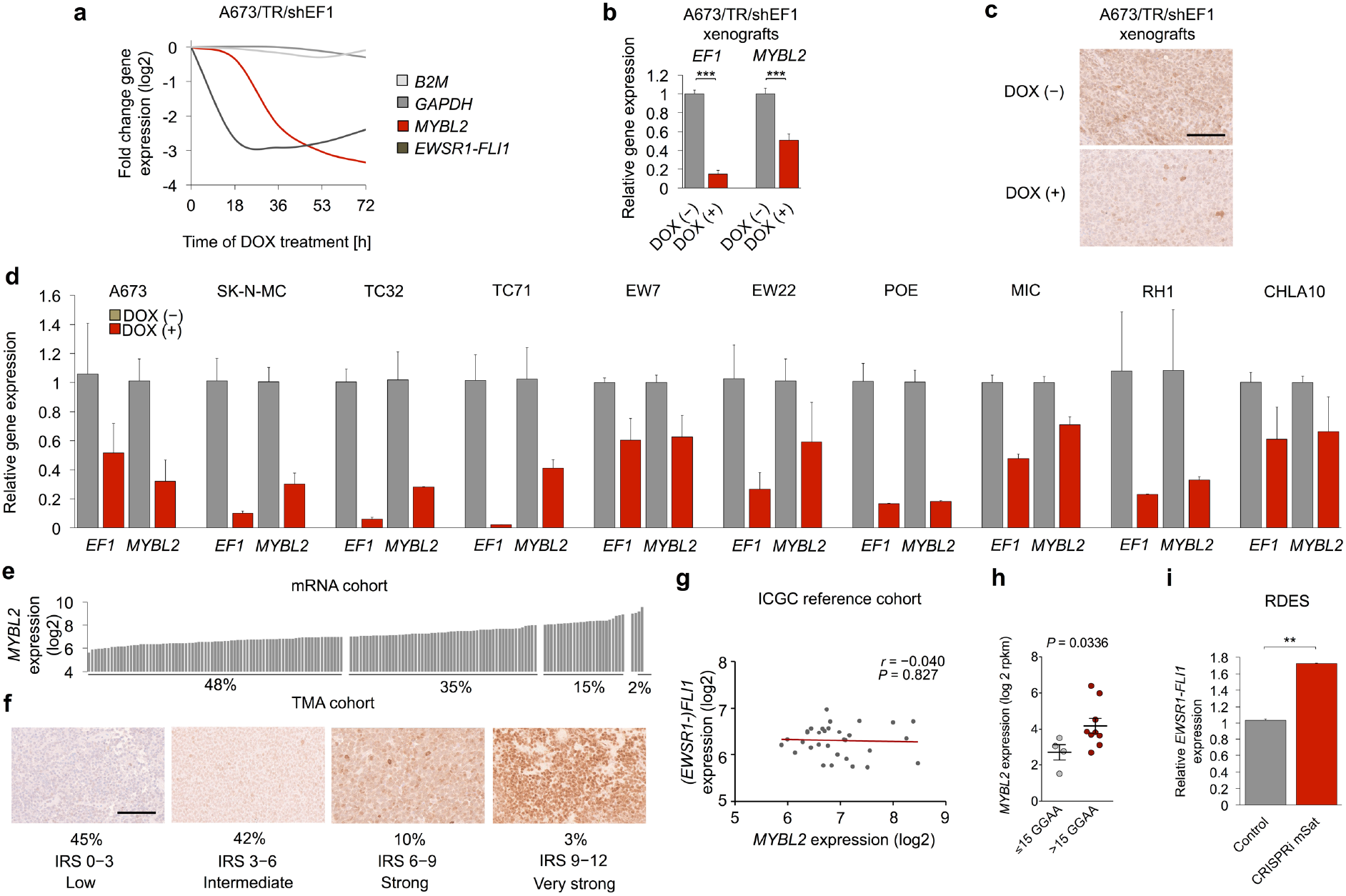
**a**, *EWSR1-FLI1, MYBL2, GAPDH* and *B2M* expression in A673/TR/shEF1 cells containing a DOX-inducible specific shRNA construct against EWSR1-FLI1, measured by Affymetrix HG-U133A2.0 arrays (GSE27524) at different time points after start of DOX addition to the media. **b**, *Ex vivo* analysis of *EWSR1-FLI1* and *MYBL2* expression by qRT-PCR in A673/TR/shEF1 xenografts either with or without addition of DOX to the drinking water. n=5 mice per group. Unpaired student’s t-test. **c**, Representative pictures of IHC p-MYBL2 staining of A673/TR/shEF1 xenografts in NSG mice either with or without addition of DOX to the drinking water. Scale bar is 50 μm. **d**, *EWSR1-FLI1* and *MYBL2* expression as measured by qRT-PCR in ten EwS cell lines containing a DOX-inducible specific shRNA construct directed against *EWSR1-FLI1* either with or without addition of DOX. **e,f** Ranked log2 *MYBL2* expression intensities in 166 primary EwS tumors as determined by Affymetrix gene expression microarrays (e) and representative images of IHC staining for p-MYBL2 of a TMA comprising 208 primary EwS tumors (f). Scale bar is 50 μm. Percentages of patients showing either low, intermediate, strong or very strong *MYBL2* expression are reported. **g**, Linear regression of *FLI1* expression onto *MYBL2* expression in an Affymetrix gene expression dataset comprising 32 primary EwS tumors with *EWSR1-FLI1* translocation. *FLI1* expression was used as a surrogate for *EWSR1-FLI1* expression, as wildtype *FLI1* is not expressed in EwS. **h**, *MYBL2* expression as determined by RNA-seq depending on the number of consecutive GGAA-repeats (lead/longer allele) of the *MYBL2-associated* GGAA-microsatellite in 13 primary EwS for which matched whole genome sequencing data was available. **i**, Analysis of relative *EWSR1-FLI1* expression in RDES EwS cells after CRISPRi-mediated inhibition of the *MYBL2-associated* GGAA-microsatellite. Data are mean and SEM, n=2. Unpaired student’s t-test. *** *P*<0.001, ** *P*<0.01, * *P*<0.05.

**Supplementary Figure 2:**
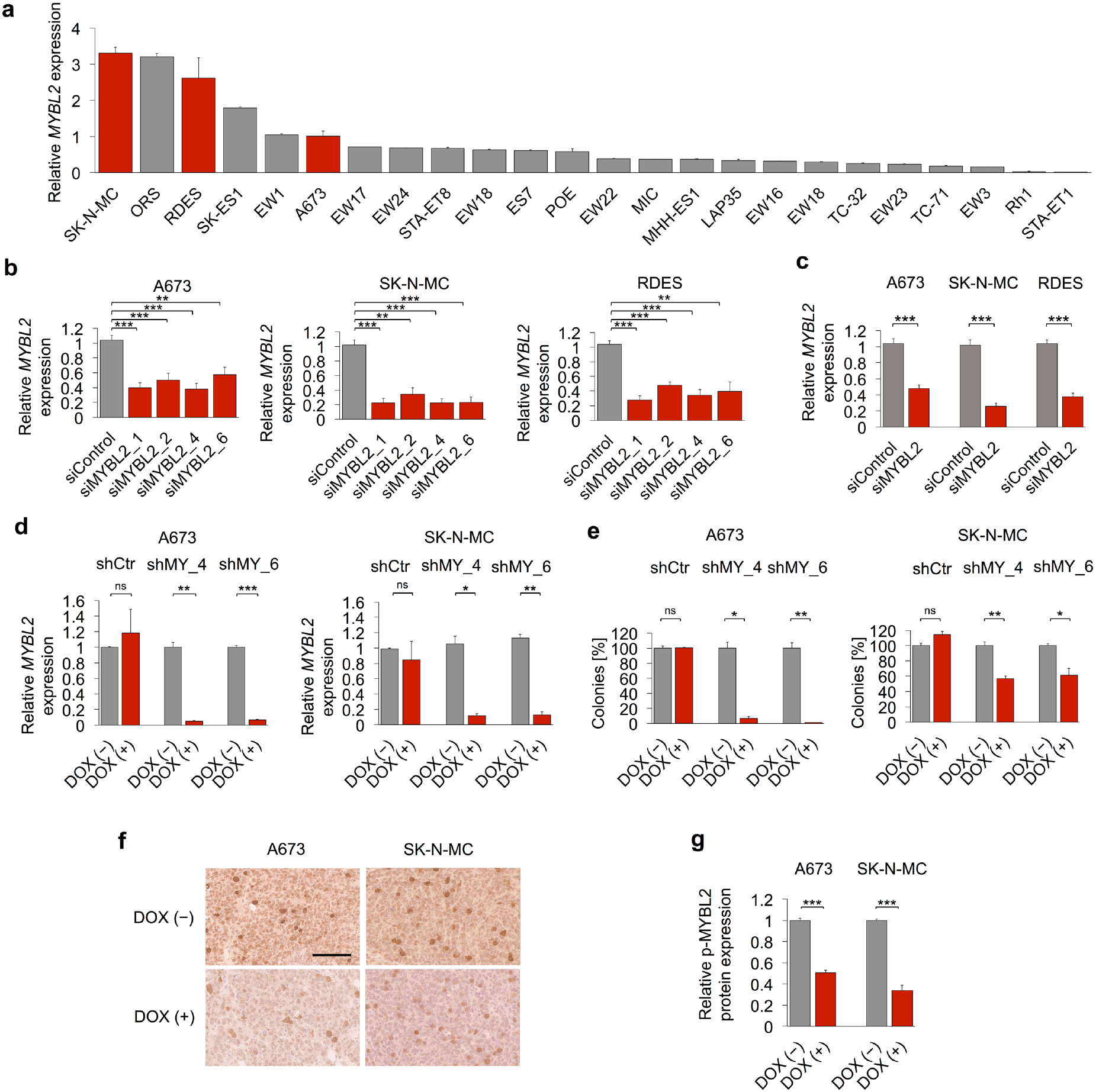
**a**, *MYBL2* expression in 24 EwS cell lines, normalized to expression in A673 as measured by qRT-PCR. **b**, Relative *MYBL2* expression as measured by qRT-PCR after transfection of A673, SK-N-MC and RDES EwS cell lines with either four different specific siRNAs directed against *MYBL2* or a non-targeting siControl. Data are mean and SEM, *n*≥3. Unpaired student’s t-test. **c**, Summary of four different specific siRNAs directed against *MYBL2* shown in (b). Data are mean and SEM, *n*≥3. Unpaired student’s t-test. **d**, Relative *MYBL2* expression as measured by qRT-PCR of A673 and SK-N-MC cells containing either DOX-inducible specific shRNA constructs directed against *MYBL2* or a non-targeting shControl. Cells were grown either with or without DOX. *n*=3. Unpaired students t-test. **e**, Relative colony number of CFAs of A673 and SK-N-MC cells containing either DOX-inducible specific shRNA constructs directed against *MYBL2* or a non-targeting shControl. Cells were grown either with or without DOX. *n*=3. Unpaired students t-test. **f**, Representative images of NSG mice xenografts of A673 and SK-N-MC cells containing a DOX-inducible specific shRNA constructs directed against *MYBL2* stained for p-MYBL2. Scale bar is 50 μm. Mice were treated either with or without addition of DOX to the drinking water. **g**, Quantification of p-MYBL2 staining as described in (f). Data are represented as mean and SEM of ten high-power fields in five tumors for each condition. Unpaired students t-test. *** *P*<0.001, ** *P*<0.01, * *P*<0.05.

**Supplementary Figure 3:**
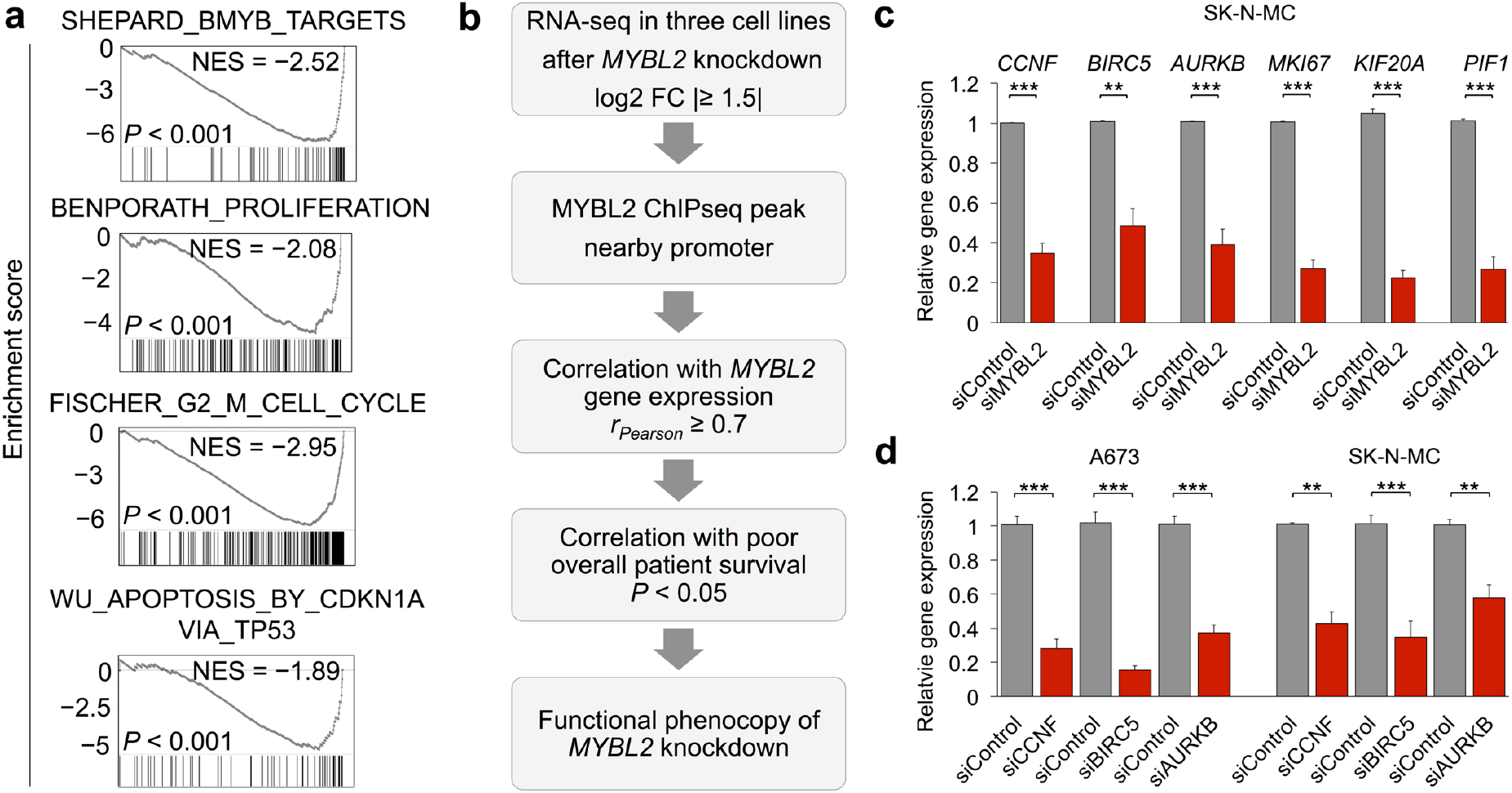
**a**, Selected negatively enriched gene sets in RNA-seq data after siRNA-mediated *MYBL2* knockdown compared to a non-targeting siControl (summary of three cell lines, *n*=3). **b**, Flow-chart showing the algorithm used to identify the top functionally relevant downstream target genes of MYBL2. **c**, Validation of downregulation of selected DEGs by qRT-PCR after *MYBL2* knockdown in SK-N-MC EwS cells, n=3. Unpaired student’s t-test. **d**, Relative gene expression of *CCNF, BIRC5* and *AURKB* expression as measured by qRT-PCR after transfection of A673 and SK-N-MC EwS cell lines with either two different specific siRNAs directed against *CCNF, BIRC5* or *AURKB* (summary of two siRNAs is shown) or a non-targeting siControl. Data are mean and SEM, n=3. Unpaired student’s t-test. *** *P*<0.001, ** *P*<0.01, * *P*<0.05.

**Supplementary Figure 4:**
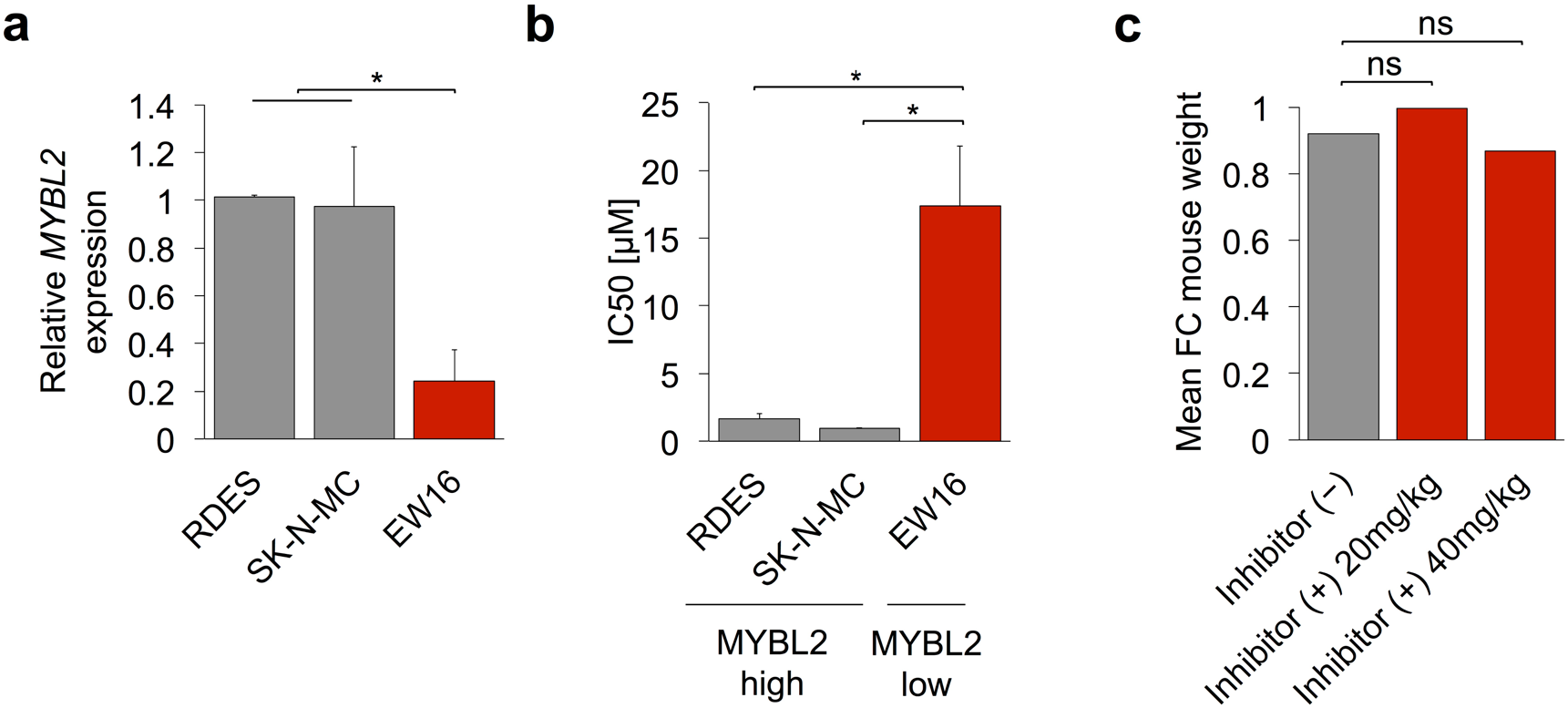
**a**, Relative *MYBL2* expression as measured by qRT-PCR of RDES, SK-N-MC and EW16 Ewing sarcoma cell lines. Data are mean and SEM, *n*=3. Unpaired student’s t-test. **b**, MYBL2 high expressing cell lines (RDES, SK-N-MC) and a MYBL2 low expressing cell line (EW16) cells were treated with the CDK2 Inhibitor NU6140 in different concentrations. Relative cell growth was assessed by Resazurin assay. Data are mean and SEM, *n*=3. Unpaired student’s t-test. **c**, Fold change (FC) of mouse weight before and after treatment with up to 40 mg/kg of the CDK2 inhibitor NU6140 for 12 days. *** *P*<0.001, ** *P*<0.01, * *P*<0.05.

